# Sirt5 regulates chondrocyte metabolism and osteoarthritis development through protein lysine malonylation

**DOI:** 10.1101/2024.07.23.604872

**Authors:** Huanhuan Liu, Anupama Binoy, Siqi Ren, Thomas C. Martino, Anna E. Miller, Craig R. G. Willis, Shivakumar R. Veerabhadraiah, Abhijit Sukul, Joanna Bons, Jacob P. Rose, Birgit Schilling, Michael J. Jurynec, Shouan Zhu

**Affiliations:** Department of Biomedical Sciences, Heritage College of Osteopathic Medicine (HCOM), Ohio University, Athens, OH, 45701, USA; Ohio Musculoskeletal and Neurological Institute (OMNI), Heritage College of Osteopathic Medicine (HCOM), Ohio University, Athens, OH, 45701, USA; Diabetes Institute (DI), Heritage College of Osteopathic Medicine (HCOM), Ohio University, Athens, OH, 45701, USA; School of Chemistry and Biosciences, Faculty of Life Sciences, University of Bradford, Bradford, BD7 1DP, UK; Department of Orthopaedics, University of Utah, Salt Lake City, Utah, 84108 USA; Buck Institute for Research on Aging, 8001 Redwood Boulevard, Novato, CA 94945, USA

## Abstract

**Objectives:** Chondrocyte metabolic dysfunction plays an important role in osteoarthritis (OA) development during aging and obesity. Protein post-translational modifications (PTMs) have recently emerged as an important regulator of cellular metabolism. We aim to study one type of PTM, lysine malonylation (MaK) and its regulator Sirt5 in OA development.

**Methods:** Human and mouse cartilage tissues were used to measure SIRT5 and MaK levels. Both systemic and cartilage-specific conditional knockout mouse models were subject to high-fat diet (HFD) treatment to induce obesity and OA. Proteomics analysis was performed in *Sirt5^-/-^* and WT chondrocytes. SIRT5 mutation was identified in the Utah Population Database (UPDB).

**Results:** We found that SIRT5 decreases while MAK increases in the cartilage during aging. A combination of Sirt5 deficiency and obesity exacerbates joint degeneration in a sex dependent manner in mice. We further delineate the malonylome in chondrocytes, pinpointing MaK’s predominant impact on various metabolic pathways such as carbon metabolism and glycolysis. Lastly, we identified a rare coding mutation in *SIRT5* that dominantly segregates in a family with OA. The mutation results in substitution of an evolutionally invariant phenylalanine (Phe–F) to leucine (Leu–L) (F101L) in the catalytic domain. The mutant protein results in higher MaK level and decreased expression of cartilage ECM genes and upregulation of inflammation associated genes.

**Conclusions:** We found that Sirt5 mediated MaK is an important regulator of chondrocyte cellular metabolism and dysregulation of Sirt5-MaK could be an important mechanism underlying aging and obesity associated OA development.

## INTRODUCTION

Osteoarthritis (OA) is a degenerative joint disease that affects millions of people worldwide (*1, 2*), with no currently available disease-modifying drugs. One of the central features of OA is gradual loss of articular cartilage. It has been increasingly recognized that chondrocyte metabolic dysfunctions during various OA risk conditions such as aging, obesity, and injury is an important mechanism for cartilage degeneration (*3*). However, it is still unknown how dysregulated cellular metabolism happens and how to correct it for potential therapeutics development.

Protein post-translational modifications (PTMs) are key regulators of cellular functions and metabolic pathways (*4–6*). Abnormal accumulation of these PTMs in metabolically active organs, such as the liver, has been found to play an important role in development of obesity and aging-associated diseases. For example, increased acetylation of proteins in the liver that regulate fatty acid oxidation accelerates the development of metabolic syndrome (*7*). We previously reported that there is an abnormal accumulation of protein post-translational malonylation, succinylation, and acetylation at lysine residues in the cartilage of obese mice (*8*). This presumably is due to accumulation of reactive acyl-group metabolic byproducts during obesity such as malonyl-CoA, succinyl-CoA, and acetyl-CoA, a phenomenon termed as ‘cellular carbon stress’ (*5*). Mammals possess 7 sirtuins (Sirt1-7), several of which have the ability to remove those PTMs (*4*). Sirtuin 5 (Sirt5) is a NAD^+^-dependent deacylase that can remove various acyl groups from lysine residues, such as malonyl (*9*), succinyl (*9*), and glutaryl (*10*) groups. Through its demalonylation activity, Sirt5 can modulate the levels of lysine malonylation (MaK), a PTM that has been demonstrated to mainly target glycolysis in the liver and linked to metabolic disorders (*11*). We reasoned that Sirt5-MaK signaling axis could potentially play an important role in chondrocyte functions and OA development, as chondrocytes rely substantially on glycolysis to generate ATP (*12–14*). Indeed, we previously discovered that Sirt5 expression declines in chondrocytes during aging, leading to increased MaK and altered glucose metabolism (*15*). Moreover, obesity also increases MaK in chondrocytes, which may further impair their function and contribute to OA development.

In this study, we aimed to explore the age-related changes in SIRT5-MaK signaling and investigate the impact of SIRT5 deficiency in combination with obesity on chondrocyte function and elucidate the molecular mechanisms by which SIRT5 regulates cartilage homeostasis.

Additionally, we identified a rare, dominantly inherited SIRT5 mutation (*SIRT5^F101L^*) in a familial OA cohort and examined its functional consequences in chondrocytes. Our findings provide new insights into the role of SIRT5 in cartilage metabolism and OA pathogenesis, offering potential avenues for therapeutic intervention in this debilitating disease.

## METHODS

### Animals and diet experiment

Sirt5 heterozygous knockout mice (*B6;129-Sirt5tm1FWa/J*, Stock No. 012757), *Aggrecan- Cre^ERT2^* mice (*B6.Cg-Acan^tm1(cre/ERT2)Crm^/J*, Stock No. 019148), and *Sirt5^flox/flox^* mice (*B6.129S(SJL)-Sirt5^tm1.1/cs^/AuwJ*, Stock No.033456) were purchased from The Jackson Laboratory. The mice were then bred to generate the genotype required for this study and treated with different diets. Detailed information can be found in the supplementary materials.

### Identification of human SIRT5 mutation

We used the Utah Population Database (UPDB) to identify SIRT5 mutation. The detailed diagnostic and procedure codes used to identify individuals with osteoarthritis in the UPDB can be found in the supplementary materials.

Other detailed information about the methods and materials can be found in the supplementary materials.

## RESULTS

### Sirt5 decreases while MaK increases in the cartilage during aging

To investigate the impact of aging on MaK and Sirt5 in joint tissues, we conducted immunohistochemical staining of MaK and SIRT5 in human knee articular cartilage samples obtained from donors spanning different age groups. In the young group, a majority of chondrocytes exhibited robust SIRT5 expression, whereas MaK staining was faint. Conversely, in the elderly group, MaK expression was markedly enhanced while SIRT5 staining was notably reduced (Figure 1A). Further quantitative analysis assessed the proportions of chondrocytes strongly stained, weakly stained, or unstained for SIRT5 and MaK relative to total chondrocyte counts. The results indicated a nearly 70% decline in the ratio of strongly SIRT5-stained chondrocytes in older donors compared to younger ones, whereas MaK expression showed an opposite trend (Figure 1B). To corroborate these findings in humans, we examined cartilage tissue from mice of different ages: young (25 weeks), middle-aged (48 weeks), and old (90 weeks). Consistently, we observed a significant age-associated increase in MaK levels (Figure 1C and 1D).

**Figure 1.**
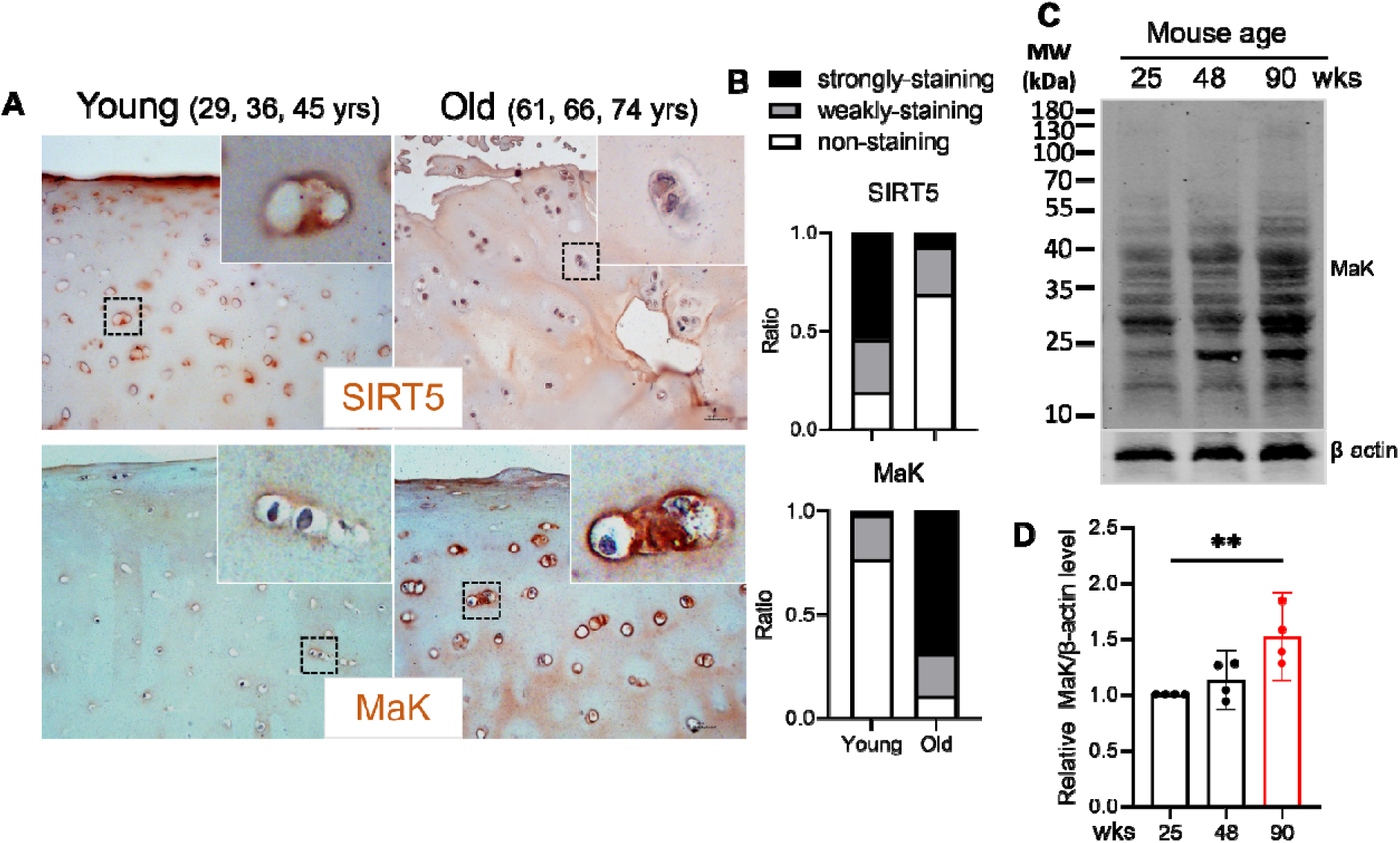
Sirt5 decreases while MaK increases in cartilage during aging. (**A**) Immunohistochemical staining for SIRT5 and lysine malonylation (MaK) in the human articular cartilage tissues from young and old donors (n=3). Insert boxes show the magnified images of the dashed squared area. (**B**) Quantification of the ratio of strongly-, or weakly-, or non-stained chondrocytes to total chondrocytes. (**C**) Immunoblotting for MaK and β-actin in the articular cartilage tissues from mice of different ages (25, 48, and 90 wks). (D) Quantification of the MaK relative to β-actin level (n=4). Data are expressed as the means ± SD, \**P* < 0.05, \*\**P* < 0.01, and \*\*\**P* < 0.001 by Student’s *t* test.

In summary, our findings suggest that aging is associated with altered expression patterns of MaK and SIRT5 in both human and mouse joint tissues, with MaK levels increasing significantly with age. These results highlight potential age-related changes in metabolic pathways within chondrocytes that merit further investigation.

### Sirt5 systemic knockout (*Sirt5^-/-^*) exacerbates HFD induced metabolic dysfunctions and OA development in mice in a sex dependent manner

In our previous work, we highlighted Sirt5 as a critical regulator of chondrocyte cellular metabolism (*15*), noting that *Sirt5^-/-^* mice exhibit mild spontaneous cartilage degeneration by 10 months of age (*8*). We also demonstrated that another major OA risk factor, obesity, also increases MaK in the cartilage tissue (*15*). To explore the role of Sirt5 in high-fat diet (HFD)- induced osteoarthritis (OA) development, we subjected both WT and *Sirt5^-/-^*mice to 20 weeks of either HFD or low-fat diet (LFD) to induce obesity. Prior to dietary intervention, female *Sirt5^-/-^*mice exhibited lower body weight compared to female WT mice, a distinction maintained under LFD conditions but gradually diminishing under HFD (Figure 2A, left panel). Similar weight disparities were observed in males pre-diet and with LFD, though HFD induced less weight gain in *Sirt5^-/-^* mice compared to WT (Figure 2B). Notably, despite lesser weight gain, *Sirt5^-/-^* mice displayed greater glucose intolerance than WT counterparts irrespective of sex (Figure 2C and 2D), indicative of systemic metabolic dysfunctions associated with Sirt5 deficiency. Furthermore, analysis of body composition revealed that HFD resulted in reduced lean mass percentage in both WT and *Sirt5^-/-^*mice across sexes. Interestingly, HFD led to increased fat mass specifically in female *Sirt5^-/-^* mice, whereas both male WT and *Sirt5^-/-^* mice showed elevated fat mass under HFD conditions (Figure 2E). These findings underscore the complex interplay between Sirt5 and metabolic homeostasis, highlighting Sirt5 as a potential modulator of diet-induced metabolic changes.

**Figure 2.**
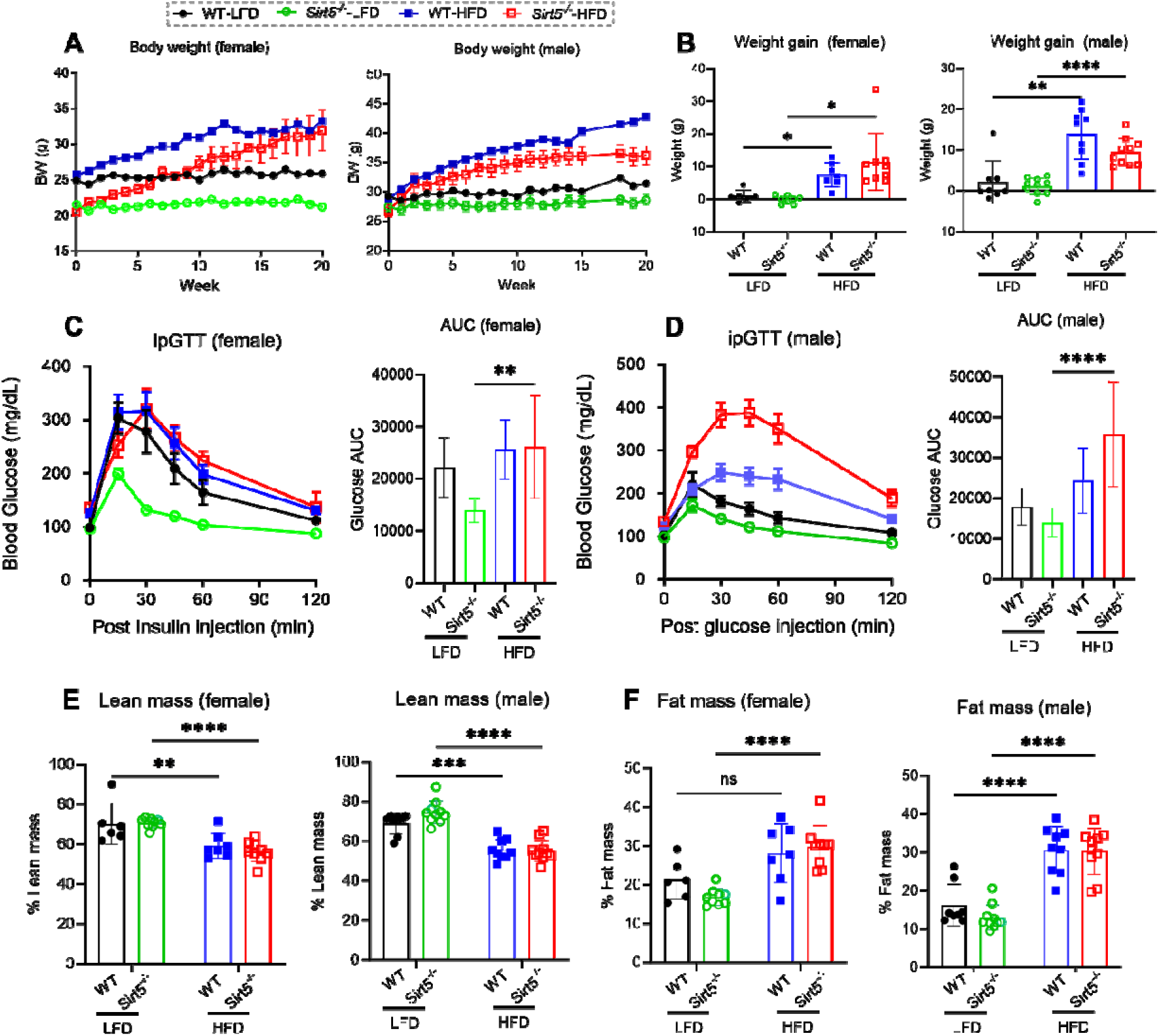
Sirt5 systemic knockout perturbs metabolism in mice. (**A**) Weight changes of *Sirt5^-/-^* and WT mice during a 20-week high-fat diet (HFD) or low-fat diet (LFD) feeding. Females (n=6 for the WT-LFD, n=7 for the WT-HFD, n=9 for the *Sirt5^-/-^*-LFD, and n=9 for the *Sirt5^-/-^*- HFD) are on the left and males (n=8 for the WT-LFD, n=9 for the WT-HFD, n=10 for the *Sirt5^-/-^*-LFD, and n=10 for the *Sirt5^-/-^*-HFD) are on the right. (**B**) Weight changes of *Sirt5^-/-^* and WT mice after the diet feeding. Females are on the left and males are on the right. (**C**) Intraperitoneal Glucose Tolerance Test (ipGTT) in female *Sirt5^-/-^*and WT mice after HFD or LFD diet feeding (left panel). The areas under the curve based on the ipGTT results are depicted in the right panel. (**D**) Intraperitoneal Glucose Tolerance Test (ipGTT) in male *Sirt5^-/-^* and WT mice after HFD or LFD diet feeding (left panel). The areas under the curve based on the ipGTT results are depicted in the right panel. (**E**) Lean mass of *Sirt5^-/-^* and WT mice after the diet feeding. Females are on the left and males are on the right. (**F**) Fat mass of *Sirt5^-/-^*and WT mice after the diet feeding. Females are on the left and males are on the right. Data are expressed as the means ± SD, \**P* < 0.05, \*\**P* < 0.01, and \*\*\**P* < 0.001 by two-way ANOVA.

Next, we evaluated the knee joints of these mice. Under LFD conditions, *Sirt5^-/-^* mice exhibited early signs of proteoglycan loss compared to WT mice (Figure 3A, left panel). With HFD treatment, WT mice showed mild to moderate cartilage degeneration with superficial lesions across the joint surface, whereas *Sirt5^-/-^* mice displayed deeper and larger cartilage defects (Figure 3A, right panel). While no statistically significant differences were observed in OARSI scores among the groups (Figure 3B and 3I), male *Sirt5^-/-^*mice notably exhibited higher Mankin scores compared to their WT counterparts (Figure 3C and 3J). HFD exacerbated these differences more prominently in males than females (Figure 3C and 3J). Conversely, no significant differences were detected in osteophyte scores (Figure 3D and 3K) or Tidemark duplication scores (Figure 3F and 3M) between genotypes or sexes. *Sirt5^-/-^*mice also exhibited significantly higher cartilage damage scores, particularly exacerbated by HFD in males (Figure 3E and 3L). Regarding Safranin-O staining, female *Sirt5^-/-^* mice showed more than twice the loss of staining intensity of WT mice in both LFD and HFD groups (Figure 3G). In males, significant differences were observed only under HFD conditions (Figure 3N). Interestingly, hypertrophic chondrocytes were more prevalent in *Sirt5^-/-^* mice under both LFD and HFD conditions, particularly pronounced in males (Figure 3H and 3O). These findings underscore the differential impact of Sirt5 deficiency on cartilage integrity and osteoarthritis progression, influenced by diet and gender-specific factors.

**Figure 3.**
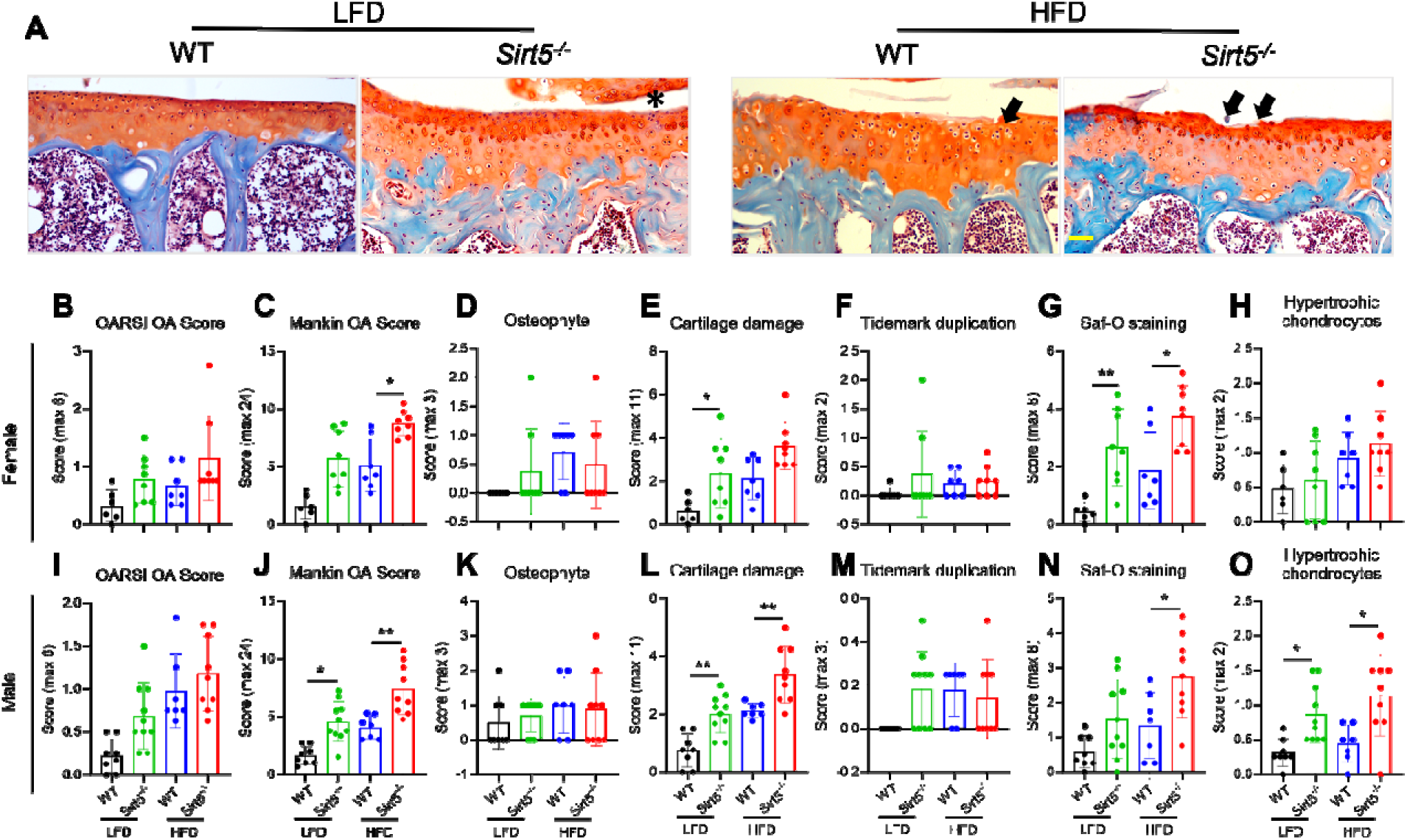
Sirt5 systemic knockout mice develop more severe HFD induced OA. (**A**) Fast Green/Safranin Orange staining of *Sirt5^-/-^* and WT mouse knee joints (28 weeks old). Pictures show the medial tibias. Asterix indicates loss of proteoglycan staining and superficial degeneration. Arrows indicate cartilage defects. (**B-H**) Quantifications of OA severity in female *Sirt5^-/-^*and WT mouse knee joints. The OA scores include OARSI (**B**), Mankin (**C**), Osteophyte (**D**), Cartilage damage (**E**), Tidemark duplication (**F**), Saf-O staining (**G**), and Hypertrophic chondrocytes (**H**). (**H-O**) Quantifications of OA severity in male *Sirt5^-/-^*and WT mouse knee joints. . Data are expressed as the means ± SD, \**P* < 0.05, \*\**P* < 0.01, and \*\*\**P* < 0.001 by two- way ANOVA.

### Sirt5 cartilage specific knockout (*Sirt5-CKO*) does not affect systemic metabolism but promotes HFD induced OA development in mice in a sex-dependent manner

To elucidate the specific role of Sirt5 in joint-specific osteoarthritis (OA) development, we utilized *Aggrecan-Cre^ERT2^* mice crossed with *Sirt5^flox/flox^* mice to achieve cartilage-specific knockout of Sirt5. Similarly, both wild-type (WT; *Sirt5^flox/flox^*) and *Sirt5* conditional knockout (*Sirt5-CKO; Aggrecan-Cre^ERT2^;Sirt5^flox/flox^*) mice were administered Tamoxifen to induce *Sirt5* deletion, followed by exposure to either a HFD or LFD to induce obesity. Unlike systemic knockout models, WT and *Sirt5-CKO* mice showed similar weight gain under HFD conditions in both males and females (Figure 4A and 4B). Additionally, both groups exhibited comparable glucose intolerance following HFD treatment, as evidenced by nearly twofold increases in the area under the curve (AUC) during the intraperitoneal glucose tolerance test (ipGTT) compared to LFD groups, irrespective of sex (Figure 4C and 4D). Furthermore, HFD led to a reduction in lean mass in both WT and *Sirt5-CKO* mice regardless of sex (Figure 4E). Interestingly, only WT female mice, but not *Sirt5-CKO* females, showed a significantly higher percentage of fat mass under HFD conditions, possibly due to greater variability in fat accumulation among *Sirt5-CKO* mice. Conversely, in males, both WT and *Sirt5-CKO* mice exhibited significantly increased fat mass percentages after HFD exposure (Figure 4F). These findings highlight the joint-specific implications of Sirt5 in metabolic responses to diet-induced obesity, independent of its effects on systemic metabolism.

**Figure 4.**
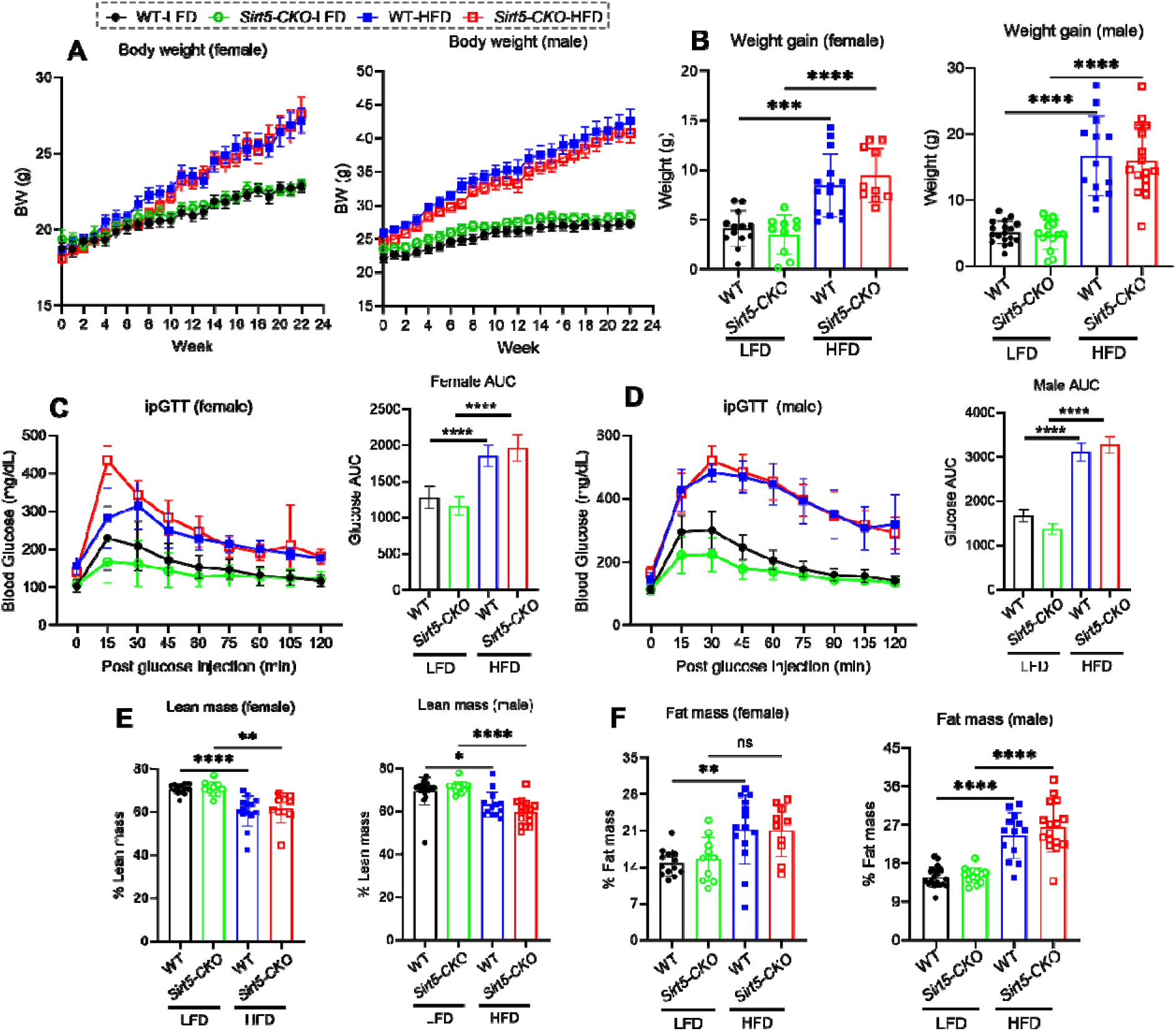
Sirt5 cartilage specific knockout (*Sirt5-CKO*) minimally perturbs metabolism in mice. (**A**) Weight changes of *Sirt5-CKO* and WT mice during a 24-week HFD or LFD feeding. Females (n=6 for the WT-LFD, n=7 for the WT-HFD, n=9 for the *Sirt5^-/-^*-LFD, and n=9 for the *Sirt5^-/-^*-HFD) are on the left and males (n=13 for the WT-LFD, n=10 for the WT-HFD, n=14 for the *Sirt5^-/-^*-LFD, and n=10 for the *Sirt5^-/-^*-HFD) are on the right. (**B**) Weight changes of *Sirt5- CKO* and WT mice after the diet feeding. Females are on the left and males are on the right. (**C**) Intraperitoneal Glucose Tolerance Test (ipGTT) in female *Sirt5-CKO* and WT mice after HFD or LFD diet feeding (left panel). The areas under the curve based on the ipGTT results are depicted in the right panel. (**D**) Intraperitoneal Glucose Tolerance Test (ipGTT) in male *Sirt5-CKO* and WT mice after HFD or LFD diet feeding (left panel). The areas under the curve based on the ipGTT results are depicted in the right panel. (**E**) Lean mass of *Sirt5-CKO* and WT mice after the diet feeding. Females are on the left and males are on the right. (**F**) Fat mass of *Sirt5-CKO* and WT mice after the diet feeding. Females are on the left and males are on the right. Data are expressed as the means ± SD, \**P* < 0.05, \*\**P* < 0.01, and \*\*\**P* < 0.001 by two-way ANOVA.

We then investigated the role of cartilage-specific Sirt5 knockout in osteoarthritis (OA) development. Initially, we confirmed efficient deletion of Sirt5 specifically in cartilage, with no impact on other tissues such as muscle (Figure 5A). Subsequent joint analysis revealed that HFD induced typical moderate superficial cartilage lesions, proteoglycan loss, and osteophyte formation in WT mice (Figure 5B). In contrast, *Sirt5-CKO* mice subjected to HFD exhibited more extensive cartilage defects covering approximately half to two-thirds of the articular surface, along with pronounced osteophyte development at the medial tibial side (Figure 5B). Similar to findings from systemic knockout studies, OARSI scores did not show statistically significant differences between groups (Figure 5C and 5K). However, under HFD conditions, *Sirt5-CKO* males displayed significantly higher Mankin scores compared to WT mice, indicative of more severe OA in *Sirt5-CKO* mice (Figure 5D and 5L), consistent with histological observations (Figure 5B). While no statistical significance was observed for osteophyte scores (Figure 5E and 5M), *Sirt5-CKO* mice exhibited higher scores, suggesting a trend towards increased osteophyte formation compared to WT. Additionally, *Sirt5-CKO* males showed higher cartilage damage scores under HFD conditions, with no differences in tidemark duplication or hypertrophic chondrocyte scores observed in either sex (Figure 5G, 5I, 5O, and 5Q). In terms of proteoglycan staining, *Sirt5-CKO* males exhibited reduced staining compared to WT males (Figure 5H and 5N).

**Figure 5.**
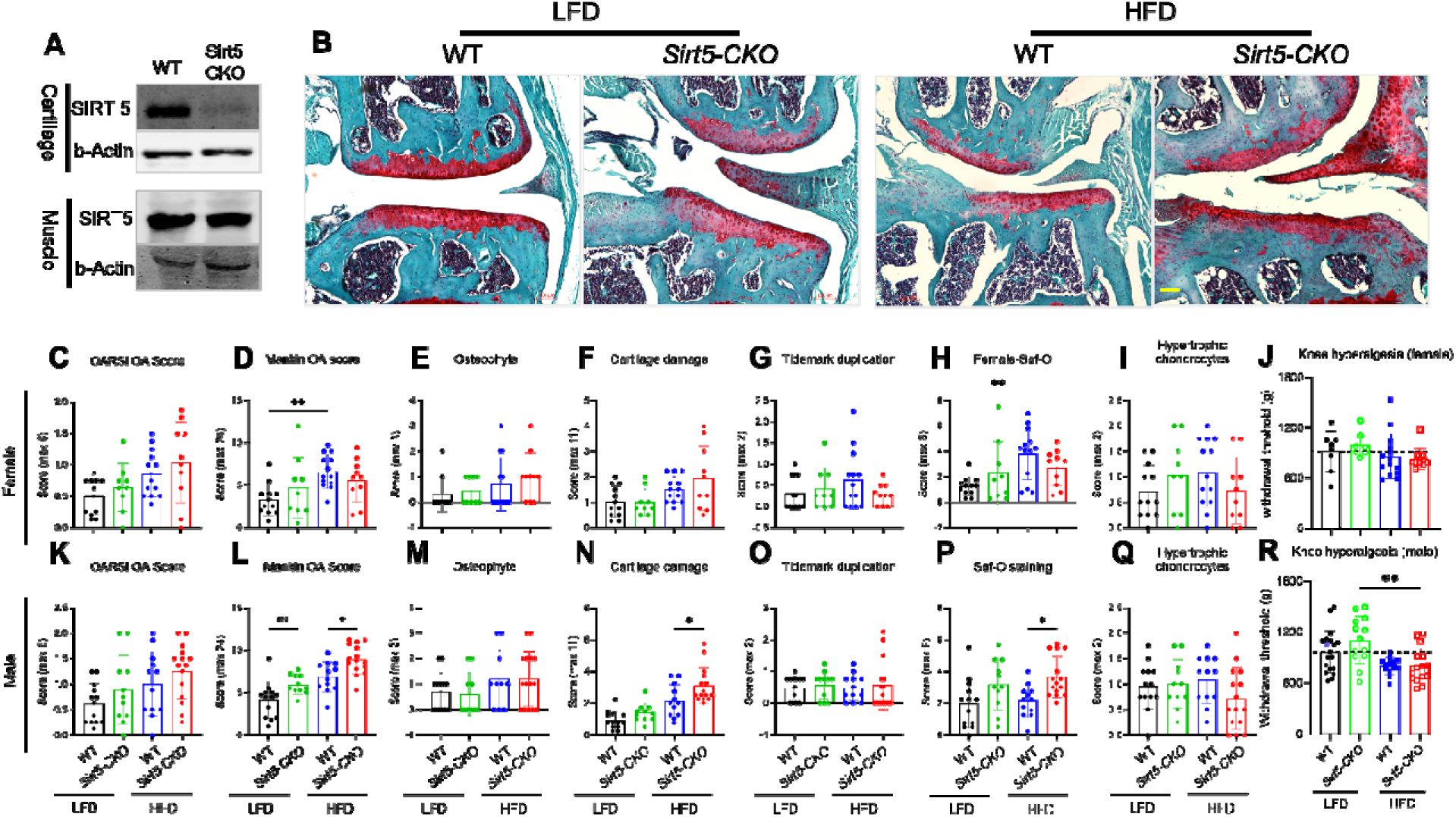
Sirt5 cartilage specific knockout mice (*Sirt5-CKO*) develop more severe HFD induced OA. (**A**) Western blotting for SIRT5 protein in both cartilage and muscle tissue in *Sirt5- CKO* and WT mice. (**B**) Fast Green/Safranin Orange staining of *Sirt5-CKO* and WT mouse knee joints (32 weeks old). Pictures show the medial compartments including both femur and tibia. (**C-I**) Quantifications of OA severity in female *Sirt5-CKO* and WT mouse knee joints. The OA scores include OARSI (**C**), Mankin (**D**), Osteophyte (**E**), Cartilage damage (**F**), Tidemark duplication (**G**), Saf-O staining (**H**), and Hypertrophic chondrocytes (**I**). (**J**) Knee hyperalgesia as an indicator for OA pain assessed in the same female mice. (**K-Q**) Quantifications of OA severity in male *Sirt5-CKO* and WT mouse knee joints. (**R**) Knee hyperalgesia as an indicator for OA pain assessed in the same male mice. Data are expressed as the means ± SD, \**P* < 0.05, \*\**P* < 0.01, and \*\*\**P* < 0.001 by two-way ANOVA.

Furthermore, we conducted a knee hyperalgesia assay using pressure application measurement (PAM) to assess joint pain. Interestingly, we observed no significant changes in knee withdrawal thresholds among female mice across different groups (Figure 5J). In contrast, *Sirt5-CKO* males under HFD conditions showed a significant reduction in knee withdrawal threshold compared to LFD, suggesting increased susceptibility to knee hyperalgesia in *Sirt5-CKO* males under obesity conditions compared to WT (Figure 5R). These results highlight the specific impact of cartilage- specific Sirt5 deficiency on OA progression and pain sensitivity in a sex-dependent manner.

### Sirt5 deficiency in chondrocytes results in decreased expression of extracellular matrix (ECM) proteins and increased expression of proinflammatory proteins

To investigate the role of Sirt5 deficiency in chondrocyte functions, we used proteomics to profile global protein expressions in WT and *Sirt5^-/-^*chondrocytes. Principle component analysis (PCA) showed a non-overlapping distribution of WT and *Sirt5^-/-^* samples with 70.7% of the variation can be explained by the principle component 1 (genotype) (Figure 6A), suggesting distinct proteomic profiles of WT and *Sirt5^-/-^* chondrocytes. Furthermore, differential abundance analysis further highlighted significant alterations, revealing 520 upregulated and 566 downregulated proteins in *Sirt5^-/-^*chondrocytes compared to WT (Figure 6B). Notably, key extracellular matrix (ECM) proteins like COL2a1 and COL11a2 were markedly downregulated, whereas inflammation-associated mediators such as prostaglandin endoperoxide synthase 1 (PTGS1) and 15-hydroxyprostaglandin dehydrogenase (HPGD) were prominently upregulated in *Sirt5^-/-^* chondrocytes (Figure 6B). To delve deeper into these changes, we performed cluster analyses using the Online Search Tool for Retrieval of Interacting Genes/Proteins (STRING). The upregulated proteins indicated involvement in crucial signaling pathways such as the MAPK cascade, integrin-mediated signaling, and actin cytoskeleton organization. Conversely, the downregulated proteins were associated with metabolic pathways and collagen formation/cartilage development processes (Figure 6C). These findings align with the established roles of Sirt5 in cellular metabolism and strongly support our hypothesis that Sirt5 plays a pivotal role in cartilage tissue homeostasis and potentially in the development of osteoarthritis (OA).

**Figure 6.**
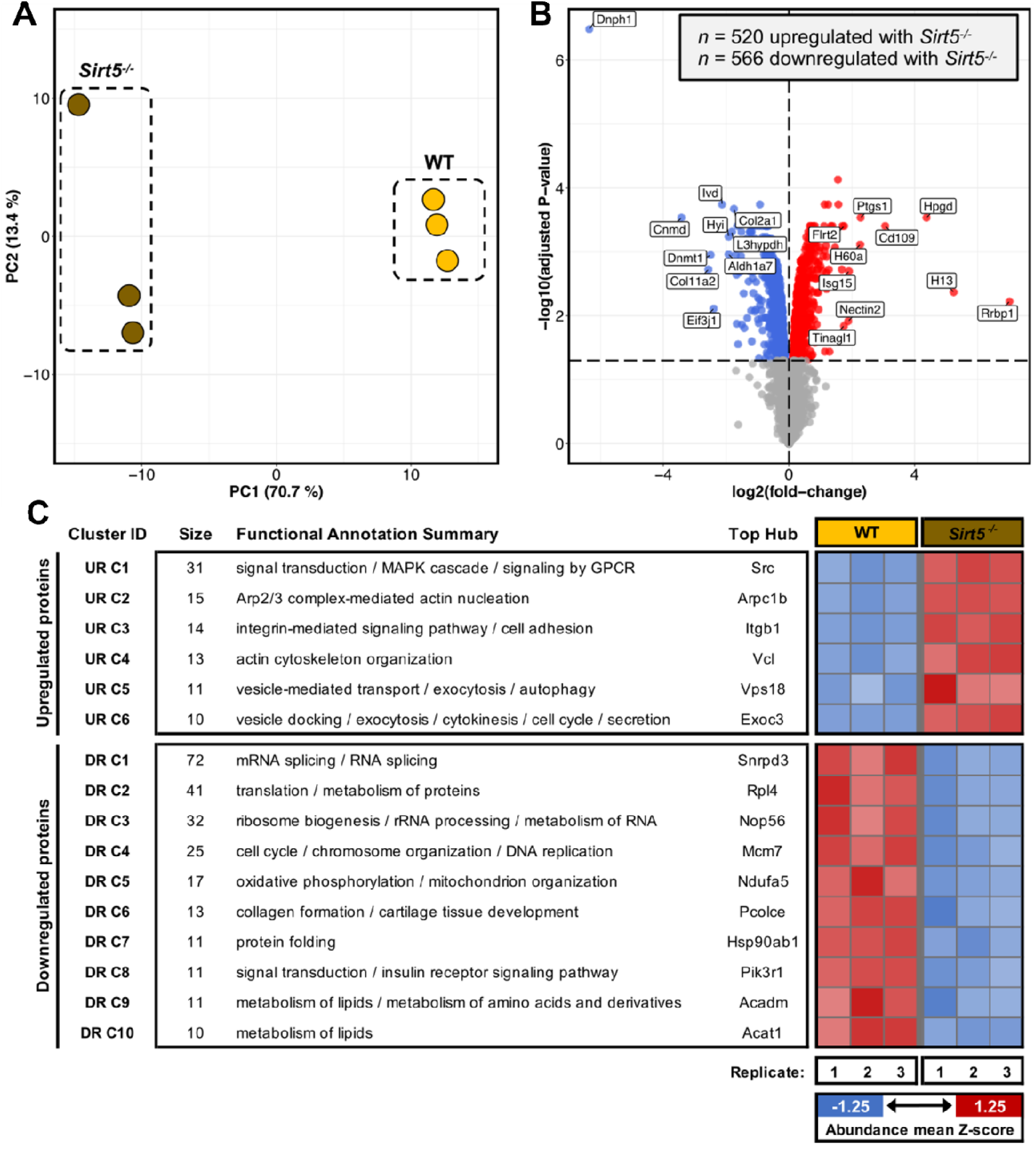
Sirt5 deficiency in chondrocytes results in decreased expression of extracellular proteins and increased expression of pro-inflammatory proteins. (**A**) Principal Component Analysis (PCA) plot of *Sirt5^-/-^* and WT mouse chondrocyte proteomic samples (n=3). (**B**) Volcano plot depicting protein differential regulation between *Sirt5^-/^*^-^ vs. WT samples. Red and blue denote significant upregulation and significant downregulation, respectively. Top 10 upregulated/downregulated proteins as based on magnitude of log2 fold-change are labelled in-plot. (**C**) Overview of protein interaction analysis. Given are all clusters of significantly upregulated/downregulated proteins that contain at least 10 proteins. Functional annotation summaries are representative based on enriched Gene Ontology biological process and Reactome pathway terms. Top hubs have been defined on eigenvector centrality. Accompanying heatmap depicts mean abundance Z-score of each cluster in each sample.

### Sirt5 regulated MaK is mainly targeting metabolic enzymes in chondrocytes

Our previous studies have demonstrated that Sirt5 plays a crucial role in regulating chondrocyte cellular metabolism and the post-translational modification of lysine residues through malonylation (*15*). To investigate the extent of malonylation in primary chondrocytes from WT and *Sirt5^-/-^* mice, we employed a previously established workflow for antibody-based enrichment and identification of peptides from protein digests containing malonylation modifications (MaK) (Figure 7A). In total, we quantified over 1,000 malonylated peptides covering 469 protein sites across all samples. Among these sites, 152 exhibited a measurable difference between WT and *Sirt5^-/-^*chondrocytes. Notably, 52.3% of the identified proteins had a single malonylation site, while 47.7% had two or more MaK sites (Figure 7B).

**Figure 7.**
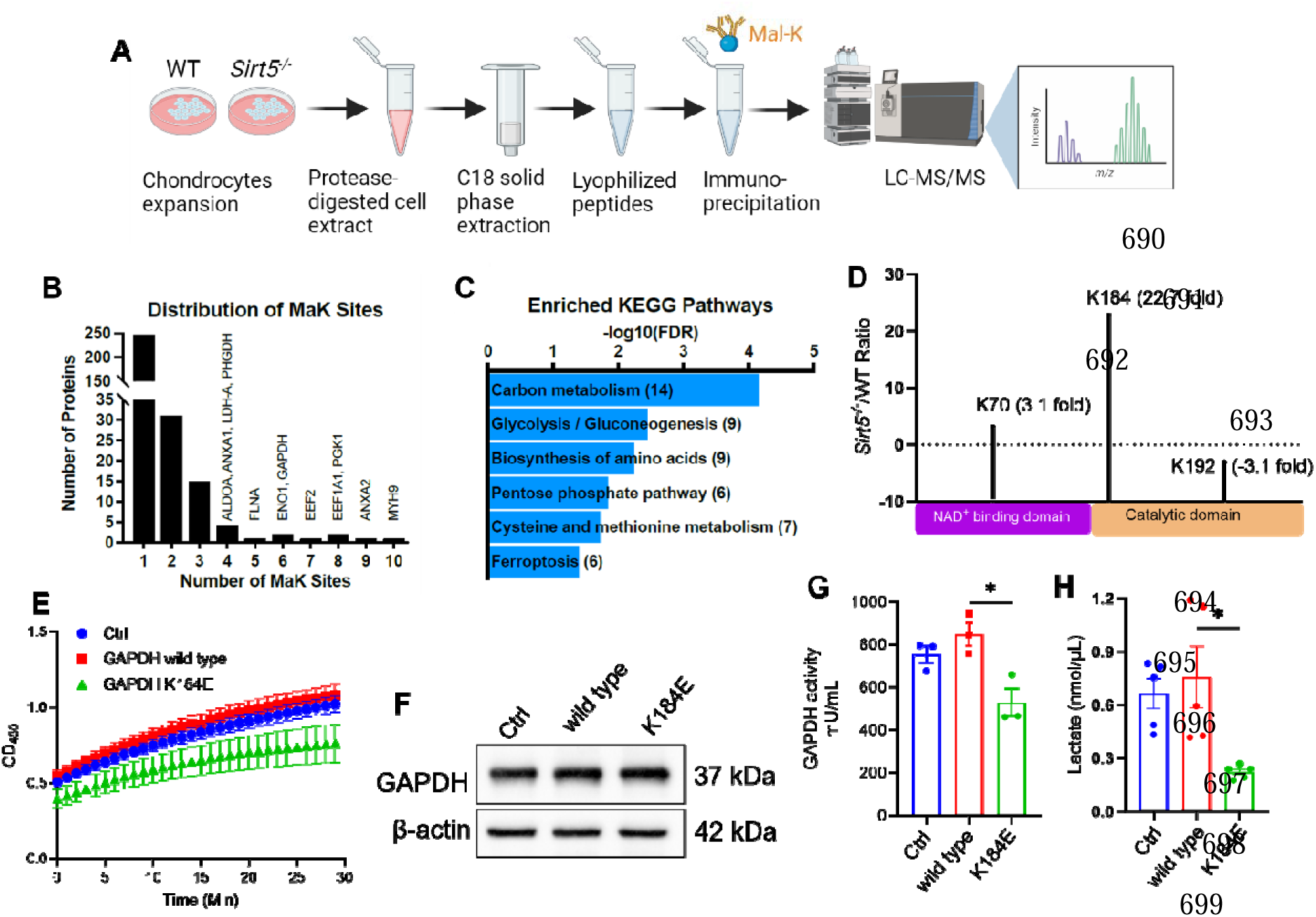
Enrichment and identification of lysine malonylome in mouse primary chondrocytes by label-free quantification. (**A**) Schematic description of experimental procedures of the label-free quantification of lysine malonylome. (**B**) Distribution of the number of MaK sites identified per protein in chondrocytes. Most malonylated proteins contain a single site, but some contain multiple sites. (**C**) KEGG pathway enrichment analysis of malonylated proteins in chondrocytes. Numbers in the brackets show the number of malonylated proteins in the pathways. (**D**) Sites of malonylation in the GAPDH protein with respect to its two functional domains, NAD binding, and catalytic domains. The ratio of malonylation between WT and *Sirt5^-/-^*samples are shown above. (**E**) GAPDH enzymatic activity kinetics assay. The assay includes on-transfected control (Ctrl), wild type GAPDH, and mutant GAPDH (K184E). (**F**) Western blotting for GAPDH to confirm successful transfection of WT and mutant GAPDH in chondrocytes. (**G**) Quantifications of GAPDH activity in mouse primary chondrocytes transfected with WT or mutant GAPDH. (**I**) Quantification of lactate production in mouse primary chondrocytes transfected with WT or mutant GAPDH. Data are expressed as the means ± SD, \**P* < 0.05, \*\**P* < 0.01, and \*\*\**P* < 0.001 by T-test.

We performed KEGG pathway enrichment analysis on the malonylated proteins and found significant enrichment in pathways related to carbon metabolism and glycolysis/gluconeogenesis (Figure 7C). Glycolysis is a fundamental metabolic pathway in chondrocytes, providing ATP, and our findings suggest that Sirt5-regulated malonylation targets key metabolic proteins in this context.

Given the enrichment of MaK modifications in glycolytic enzymes and the critical role of glycolysis in chondrocyte energy metabolism, we compared the relative abundance of MaK in all detected glycolytic enzymes between *Sirt5^-/-^* and WT chondrocytes. Among them, Glyceraldehyde 3-phosphate dehydrogenase (GAPDH), which catalyzes the conversion of glyceraldehyde 3-phosphate to glycerate 1,3-bisphosphate, showed significant dynamic regulation (with *Sirt5^-/-^*/WT ratios of 3.1, 22.7, and -3.1 for three lysine sites) (Figure 7D).

To explore the functional implications of malonylation on these glycolytic enzymes and their regulation by Sirt5, we investigated whether modification of the most malonylated lysine 184 affects GAPDH enzymatic activity. We generated a mutated mouse GAPDH expression vector in which lysine 184 was substituted with glutamate (K184E), mimicking MaK modification by introducing a negatively charged residue. Mouse primary chondrocytes were transfected with either WT or mutant GAPDH. A kinetic enzymatic assay revealed that the GAPDH K184E mutant significantly reduced enzymatic reaction rates compared to both WT GAPDH and non-transfected controls (Figure 7E, 7F, and 7G). Furthermore, we measured the final glycolysis product lactate and found that the GAPDH K184E mutant also significantly decreased lactate production compared to WT GAPDH and control groups (Figure 7G). These results underscore the functional impact of Sirt5-regulated malonylation on glycolytic enzymes in chondrocytes, highlighting a novel regulatory mechanism in cellular metabolism.

### Identification and functional analysis of a human OA-associated *SIRT5^F101L^* mutation

We further explored the clinical relevance of our study. Using the Utah Population Database (UPDB), we identified families in which osteoarthritis (OA) segregates as a dominant trait, with multiple members diagnosed with early-onset or severe OA in various joints, including the 1st metatarsophalangeal joint, erosive hand OA, finger interphalangeal joint OA, thumb OA, or shoulder OA (*16–18*). We completed genomic analyses for 151 families and identified a rare *SIRT5* coding mutation (rs201979175, c.C303G:p.F101L, minor allele frequency - 0.00004303) in one family that segregates dominantly with finger interphalangeal joint OA (hand OA). Members of this family also have OA in other joints, including the spine, wrist, and shoulder (Figure 8A). The *SIRT5^F101L^* mutation occurs in an evolutionarily conserved amino acid; the wild-type allele, phenylalanine (F), is invariant in the vertebrate lineage (Figure 8B). The F101L mutation is located in the substrate binding domain (Figure 8B) and is predicted to be damaging or disease-causing by multiple in silico pathogenicity prediction programs (e.g., SIFT, MutationTaster). These data indicate that a rare, dominant coding mutation in *SIRT5* is associated with familial OA.

**Figure 8.**
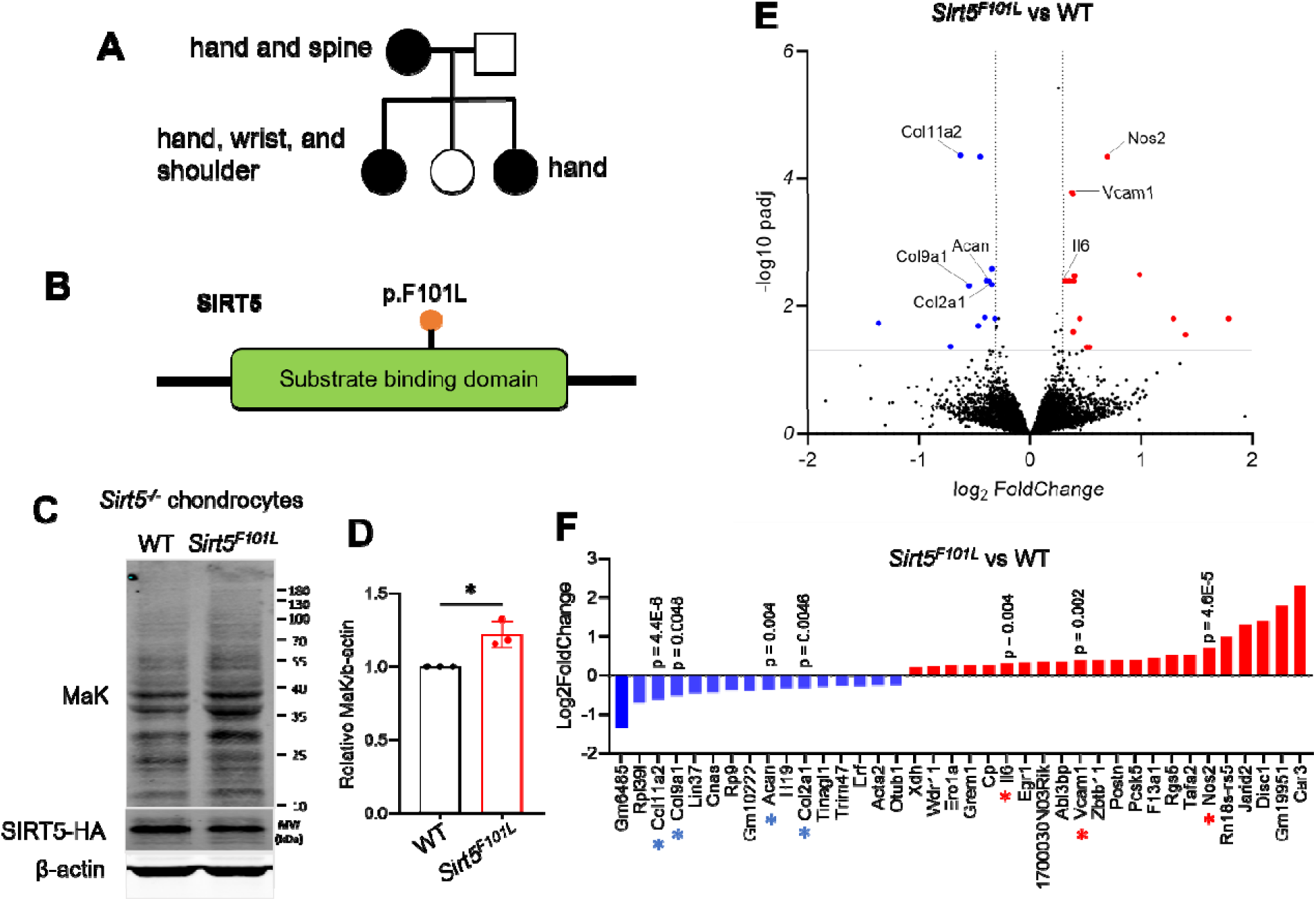
Identification of the human OA-associated *SIRT5^F101L^* mutation. (**A**) A rare *SIRT5* coding mutation dominantly segregates in a family with hand OA. Circles = females, square = males, black circles/squares = individuals affected with hand OA and individuals also had other joints affected. (**B**) Whole exome sequencing identified a mutation in a highly conserved amino acid (F101L) (F is invariant in the vertebrate lineage) in the substrate binding domain of SIRT5. (**C**) Western blotting analysis indicates chondrocytes transfected with *SIRT5^F101L^*mutant have a higher level of MaK comparing with WT. (**D**) Quantification of the band intensity between WT and *SIRT5^F101L^* mutant in the blot in panel (C). (n=3) (**D**) Bulk RNA-Seq analysis of primary mouse chondrocytes transfected with WT or *SIRT5^F101L^* mutant constructs. Expression of *Sirt5^F101L^* causes a reduction of multiple cartilage ECM genes (*Col11a2, Col9a1, Acan, and Col2a1*) and an increase of inflammation associated genes (*Nos2, Il6, and Vcam1*) (n=3). (**E**) List of upregulated or downregulated genes as colored in the panel (C). Asterix indicates statistically significantly changed genes, p values are shown above each gene. Data are expressed as the means ± SD, \**P* < 0.05, \*\**P* < 0.01, and \*\*\**P* < 0.001 by T-test.

To determine if the *Sirt5^F101L^* mutation alters Sirt5 protein function, we tested if overexpression of *Sirt5^F101L^* in primary mouse chondrocytes affected protein MaK and gene expression. Primary mouse *Sirt5^-/-^* chondrocytes were transfected with WT or *Sirt5^F101L^* and protein extracts were collected for MaK analysis. Immunoblot analysis indicates chondrocytes transfected with *Sirt5^F101L^* have a higher level of MaK compared with WT (Figure 8C and 8D). WT and *Sirt5^F101L^* transfected primary chondrocytes were also used for bulk RNA-seq analysis. Expression of *Sirt5^F101L^* caused a reduction of multiple cartilage ECM genes (*Col11a2, Col9a1, Acan,* and *Col2a1*) and an increase of inflammation associated genes (*Nos2, Il6,* and *Vcam1*) (Figure 8F). These data are consistent with changes we observe in MaK levels and protein expression in *Sirt5^-/-^* chondrocytes (Figure 2B), suggesting that *Sirt5^F101L^* is hypomorphic. In sum, our molecular, cellular and genetic data support a central role for SIRT5 in maintaining chondrocytes homeostasis.

## DISCUSSION

Our study delves into the role of Sirtuin 5 (SIRT5) and its associated post-translational lysine malonylation in chondrocyte metabolism and osteoarthritis (OA) pathogenesis. We observed that aging and obesity lead to significant alterations in the expression and activity of SIRT5, contributing to cartilage degeneration and OA development. Our findings underscore the potential of targeting SIRT5 and its regulatory pathways as a therapeutic strategy for OA.

The sirtuin family of HDACs has seven highly conserved NAD-dependent deacetylases, which were discovered in 1999/2000 (*19, 20*). Sirtuins regulate key biological processes in mammals, including many aspects of metabolism, stress response, and signaling (*21*). In addition to their well-studied roles as lysine deacetylases, certain sirtuins can also remove other types of acyl modifications from lysine residues, including propionyl, butyryl, malonyl, succinyl, and the long-chain fatty-acid-derived myristoyl and palmitoyl groups (*9, 10, 22–24*). Sirtuins regulate numerous processes *in vivo*, including metabolism, DNA repair, metastasis, apoptosis, and translation, while promoting longevity and protecting against cancer (*21*). Sirt5 was initially reported as a lysine deacetylase found in both mitochondria and cytoplasm (*25*). However, recent studies have shown it has low deacetylase activity and strong demalonylase and desuccinylase activity (*9*). Sirt5 deficiency in liver causes hypermalonylation of proteins that are enriched in glycolytic pathway, as identified by mass spectrometry of immunoprecipitated malonyl-lysine peptides (*11*). Consistently, *Sirt5^-/-^*hepatocytes show decreased production of lactic acid and reduced glycolytic flux (*11*). We previously reported that *Sirt5^-/-^*primary chondrocytes isolated *in vitro* have upregulated MaK and impaired cellular metabolism (*26*). In this study, we further discovered that SIRT5 decreases in accordance with an increase of MaK during aging in both human and mouse cartilage. MaK has also been strongly implicated in the pathogenesis of metabolic diseases. For example, a large-scale analysis of protein malonylation in livers of *db/db* mice showed upregulated lysine malonylation of metabolic enzymes involved in glucose catabolism and FA oxidation (*27*). We also previously discovered that MaK level is increased in the cartilage of obese mice compared to lean mice (*8*). These early findings prompted us to further investigate the combined role of Sirt5 deficiency and obesity in OA development. Using both systemic and conditional knockout mouse models, our results demonstrated that obesity and Sirt5 deficiency synergistically promote chondrocyte metabolic dysfunction and cartilage degeneration. This finding potentially reveals a common signaling pathway through which aging and obesity—the two major risk factors for OA—converge to promote chondrocyte metabolic dysfunction and OA development.

Emerging evidence has suggested that acyl-based PTMs at lysine residues are crucial regulators of cellular metabolism. Proteomic studies identifying the malonylome, succinylome, and acetylome have found that metabolic proteins are enriched in these datasets (*10, 11, 28–31*). Gene ontology analyses on acetylated proteins consistently reveal that metabolic pathways, including fatty acid metabolism, glycolysis, and amino acid metabolism, are among the top targets for these PTMs. However, it remains largely unknown how sites of malonylation, succinylation, and glutarylation identified in proteomic studies affect the activity of their targets and the pathophysiology at the whole-organismal level. Malonylation is a relatively unexplored PTM compared to succinylation and acetylation. Previous research has discovered that malonylation suppresses voltage-dependent anion channel 2 (VDAC2) (*32*), mTOR (*33*), glyceraldehyde 3-phosphate dehydrogenase (GAPDH) (*11*), while activating acetyl-coenzyme A (CoA) carboxylase 1 (ACC1) (*34*). Malonylation has been implicated in several pathophysiological processes, including sepsis-induced myocardial dysfunction (*32*), angiogenesis (*33*), and hepatic steatosis (*34*). In this study, we are the first to use affinity enrichment-based proteomics to identify the malonylome in chondrocytes. We demonstrated a similar suppressive effect of GAPDH malonylation on its activity and glycolysis in chondrocytes. Combined with our data from both systemic and conditional Sirt5 knockout mouse models, our findings provide strong evidence that MaK in cartilage tissue during aging and obesity conditions could be an important underlying molecular mechanism for osteoarthritis (OA) development.

Our study provides significant insights into the sex-dependent effects of Sirt5 deficiency and HFD treatment on obesity and osteoarthritis (OA) development in mice. We found that systemic Sirt5 knockout (*Sirt5^-/-^*) female mice maintained a lower body weight than wild-type (WT) under LFD, a difference that diminished with HFD. In males, *Sirt5^-/-^* mice gained less weight than WT under HFD, indicating that Sirt5 deficiency may mitigate some HFD-induced weight gain, especially in males. Despite lesser weight gain, both male and female *Sirt5^-/-^* mice showed greater glucose intolerance compared to WT controls, highlighting metabolic dysfunctions linked to Sirt5 deficiency. Interestingly, HFD increased fat mass specifically in female *Sirt5^-/-^*mice, while both male WT and *Sirt5^-/-^* mice had elevated fat mass. This suggests that female *Sirt5^-/-^*mice may be more susceptible to HFD-induced adiposity despite their lower body weight. These findings differ from a previous study showing no significant metabolic abnormalities in *Sirt5^-/-^* mice under either diet (*35*), possibly due to different HFD treatment durations (10 vs. 20 weeks). The systemic metabolic differences between WT and *Sirt5^-/-^* were not seen in the Sirt5-CKO study. These findings underscore the importance of considering sex as a biological variable in metabolic studies and suggest potential sex-specific mechanisms by which Sirt5 influences adiposity and glucose metabolism.

Interestingly, our results showed that male *Sirt5^-/-^*mice had higher Mankin scores and more cartilage damage under HFD conditions than females. Similarly, *Sirt5-CKO* mice under HFD had more cartilage defects and higher Mankin scores compared to WT mice, with males showing more severe OA. The pressure application measurement (PAM) assay demonstrated sex-dependent differences in OA pain. While knee withdrawal thresholds did not change significantly in female mice across groups, male *Sirt5-CKO* mice under HFD had a significant reduction in knee withdrawal threshold compared to LFD. These findings indicate a greater susceptibility to diet-induced OA progression in males and a potential sex-specific response to Sirt5 deficiency in joint tissues. Our results are consistent with well documented sex dependent differences of obesity and OA pathogenesis in both clinical (*36, 37*) and preclinical studies. For example, it was found that male mice seem developing more consistent OA pathologies in response to different risk factors such as joint injury (*38, 39*) and aging (*40*). Since C57BL/6J female mice are typically more resistant to the obesogenic effects of a HFD treatment (*41, 42*), they are often excluded from obesity associated OA studies (*43*). Our results, to the best of our knowledge, are the first to report that, similar to joint injury-induced OA, obesity-associated OA appears to be more severe in males compared to females.

The identification of the human *SIRT5^F101L^* mutation in a family with dominantly inherited osteoarthritis (OA) provides valuable insights into the genetic underpinnings of OA and underscores the critical role of SIRT5 in joint health. Our findings suggest that this rare mutation may reduce SIRT5 functions and contribute to the pathogenesis of OA through alterations in chondrocyte metabolism. The *SIRT5^F101L^* mutation was identified in a multigenerational family with a high incidence of OA, affecting various joints, including the spine, wrist, and shoulder. The evolutionary conservation of the phenylalanine residue at position 101 and its predicted pathogenicity by multiple in silico tools strongly indicate the functional importance of this amino acid. The mutation’s location in the substrate-binding domain suggests a potential impact on SIRT5’s enzymatic activity, further supported by our functional assays demonstrating *Sirt5^F101L^*has decreased demalonylase activity and disrupts gene expression in chondrocytes. Many (if not all) of the previously identified OA-associated mutations are non-null, even those identified from family studies (*44–50*). By studying a hypomorphic human OA-associated allele of *Sirt5^F101L^,* we could potentially uncover novel metabolic and posttranslational effects that we could not detect using a null allele of *Sirt5*. Our on-going effort includes creating a novel mouse model harboring the *Sirt5^F101L^* mutation and examine its susceptibility to different OA risk factors and identify potential chemical modulators to increase its enzymatic activity.

Our study also is not without limitations. Firstly, the mild osteoarthritis phenotype observed in mice during aging and obesity may not fully capture the complexity of the disease’s progression in more severe cases. Additionally, the use of proteomics in *in vitro* cell culture may not accurately reflect the *in vivo* environment, potentially affecting the translatability of the results. Lastly, while the data we generated using cells transfected with *Sirt5^F101L^*mutant provides valuable insights, they may not entirely replicate the physiological conditions *in vivo*. Therefore, more research directly in the *Sirt5^F101L^*mutant mice is needed to understand the mutation’s role in osteoarthritis development.

In conclusion, our study has elucidated the critical role of Sirt5 and lysine malonylation (MaK) in chondrocyte metabolism and the pathogenesis of osteoarthritis (OA) during aging and obesity. We have demonstrated that Sirt5 deficiency, combined with obesity, significantly exacerbates joint degeneration in mice, and that MaK predominantly impacts metabolic pathways such as carbon metabolism and glycolysis. Furthermore, the discovery of a rare coding mutation in SIRT5 that segregates in a family with hand OA highlights the potential genetic contributions to OA development. These findings suggest that dysregulation of the Sirt5-MaK axis could be a pivotal mechanism underlying OA development associated with aging and obesity, offering new avenues for therapeutic interventions.

## Acknowledgements

We thank the following funding support: Hevolution Foundation AGE award AGE-008 (SZ, HL), National Institutes of Health grant R01AR081804 (SZ, HL), National Institutes of Health grant R15AR080813 (SZ, HL), Arthritis National Research Foundation, The Arthritis & Aging Grant Program (SZ), American Society for Bone and Mineral Research FIRST award (SZ), Rheumatology Research Foundation Innovative Award (SZ), Osteopathic Heritage Foundation Ralph S. Licklider, D.O. Endowed Professorship (SZ), National Institutes of Health grant R01AR082973 (MJJ), Skaggs Foundation for Research (MJJ). We thank the staff (including Tammy Mace, Angela Smith, Shawn Rosensteel, and Scott Carpenter) at the animal facility in the Life Science Building for their excellent care for our animals. We thank Daniella Issa for her contributions to the *Sirt5^-/-^*mouse colony maintenance and Shadi Moradi for her help with the tissue dissections. We acknowledge the support of instrumentation for the Orbitrap Eclipse Tribrid system from the NCRR shared instrumentation grant 1S10 OD028654 (PI: Birgit Schilling). We thank the Chicago Center on Musculoskeletal Pain (funded by NIAMS P30AR079206, PI: Anne-Marie Malfait and Rachel E. Miller) for their training on using PAM to measure knee hyperalgesia in mice.

## Author contributions

Conceptualization: SZ, HL

Methodology: HL, AB, SR, TCM, AEM, CRGW, SRV, AS, MKL, SZ

Investigation: HL, AB, SR, CRGW, MKL, SZ

Visualization: CRGW, SRV, AS, SZ

Funding acquisition: SZ, HL

Project administration: SZ, HL

Supervision: SZ

Writing – original draft: SZ

Writing – review & editing: All authors.

## SUPPLEMENTARY MATERIALS AND METHODS

### Animals and diet experiment

All animal studies were reviewed and approved by the AAALAC International accredited Institutional Animal Care and Use Committee at the Ohio University.

Sirt5 heterozygous knockout mice (*B6;129-Sirt5tm1FWa/J*, Stock No. 012757) were purchased from The Jackson Laboratory after Cryo-recovery of embryos. The mice were then bred to generate both WT and Sirt5 homozygous knockout (*Sirt5*^-/-^) mice. *Aggrecan-Cre^ERT2^* mice (*B6.Cg-Acan^tm1(cre/ERT2)Crm^/J*, Stock No. 019148) and *Sirt5^flox/flox^* mice (*B6.129S(SJL)- Sirt5^tm1.1/cs^/AuwJ*, Stock No.033456) were purchased from the Jackson Laboratory. The *Aggrecan-Cre^ERT2^* and *Sirt5^flox/flox^*were then bred together to generate WT (*Sirt5^flox/flox^*) and Sirt5- CKO (*Aggrecan-Cre^ERT2^; Sirt5^flox/flox^*) mice. The C57BL/6J mice of different ages (25, 48, 90 wks) were purchased from the Jackson Laboratory (Stock No. 000664), the cartilage of these mice was further used to extract protein and immunoblot for MaK and β-actin.

Prior to the special diet treatment, mice were group housed in a specific pathogen free facility under a controlled environment (22 ± 3°C on 14 h:10 h light/dark cycles) in ventilated cages (≤5 animals/cage) with ad libitum access to chow (Lab Diet 5053) and sterilized water. For the Sirt5 systemic knockout study, both WT and *Sirt5*^-/-^ mice were fed LFD (10% kcal fat; D12450Ji, Research Diets) or HFD (60% kcal fat; D12492i) from 8 to 28 weeks of age (n=6 for the WT-LFD, n=7 for the WT-HFD, n=9 for the *Sirt5^-/-^*-LFD, and n=9 for the *Sirt5^-/-^*-HFD). At 27 weeks of age, body fat was measured by Whole-Body Composition Analyzer (Bruker Minispec Live Mice Analyzer, LF50), and glucose tolerance testing (GTT) was performed as previously described (*1*). For the Sirt5 conditional knockout study, both WT and *Sirt5-CKO* mice at 6 weeks of age were treated with Tamoxifen (15mg/ml in 150 µL peanut oil solution) through intraperitoneal (IP) injections for 5 consecutive days. Then both WT and *Sirt5-CKO* mice were fed LFD or HFD from 8 to 32 weeks of age. Similar GTT and body composition analysis were also conducted one week before the tissue collections.

### Mouse knee hyperalgesia measurement

Mouse knee joint hyperalgesia was assessed by a Pressure Application Measurement (PAM) device (Ugo Basile, Varese, Italy) as previously described (*2*). Briefly, mice were restrained by hand and the hind paw was lightly pinned with a finger in order to hold the knee in flexion at a similar angle for each mouse. With the knee in flexion, the PAM transducer was pressed against the medial side of the ipsilateral knee while the operator’s thumb lightly held the lateral side of the knee. The PAM software guided the user to apply an increasing amount of force at a constant rate (30 g/s). If the mouse tried to withdraw its knee, the force at which this occurred was recorded. Two measurements were taken per knee and the withdrawal force data were averaged as the final data value. The operator was blinded to the genotypes of the mice.

### Immunohistochemical (IHC) staining for MAK and SIRT5 in human cartilage tissue

Human knee cartilage tissues from young (29, 36, 45 years old) and old (61, 66, 74 years old) donors were procured from either tissue banks or from patients undergoing knee arthroplasty (approved by Scripps Institutional Review Board). The tissue blocks were resected from the central weight-bearing area of the medial femoral condyle. Sections from young and old human cartilage tissues were deparaffinized, rehydrated, and incubated with antigen retrieval R-Buffer A (EMS, 62707-10) at 80 °C for 1 hour. Slides were then treated with 2% H_2_O_2_, blocked using 5% BSA, and incubated overnight at 4 °C with primary antibody anti-malonyl-lysine (Cell Signaling Technology, #14942, 1:250 dilution) or anti-SIRT5 (Cell Signaling Technology, #8782, 1:250 dilution) overnight. Negative controls were processed in parallel with rabbit IgG. Staining was detected using goat anti-rabbit antibody conjugated with HRP (Horseradish Peroxidase) with HIGHDEF Red IHC Chromagen HRP (Enzo, 11141807) for color development following manufacturer’s protocol. Sections were also counter-stained with hematoxylin for 30 seconds. The number of strongly-stained, weakly-stained, and non-stained chondrocytes were counted. The ratios of strongly-stained, weakly-stained, and non-stained chondrocytes were then calculated accordingly (ratio = # of positively stained chondrocytes/total # of chondrocytes).

### Mouse primary chondrocyte isolation and culture

Primary mouse chondrocytes were isolated from 7 day old WT and *Sirt5^-/-^*mice according to a previously published protocol(*3*). Briefly, both tibial and femoral cartilage tissue were dissected from the knee joint. The cartilage was then incubated in 3 mg/mL Collagenase D (Roche) in DMEM solution for two 45-minute periods and transferred to 0.5 mg/mL Collagenase D in DMEM solution supplemented with 3% Liberase TL (Sigma) overnight. The tissue was then homogenized by pipetting up and down to release and suspend the cells, and the homogenate was filtered through a 40 µm strainer to remove large debris. Passage zero cells were cultured and expanded in 6-well plates in complete DMEM media (Life Technology, 10567014) supplemented with 10% fetal bovine serum (FBS) and 1% penicillin/streptomycin (P/S) at 37 °C and 5% CO2. Chondrocytes reached confluency after 3-4 days in culture. Chondrocytes were then trypsinized, counted, and reseeded as passage 1 cells for specific experiments described in detail in the following sections.

### Mouse joint histology

Joint samples were collected from WT and *Sirt5^-/-^* mice (males and females with different diets) or WT and *Sirt5-CKO* mice (males and females with different diets) and used for histology. Isolated knee joints were fixed in 4% formaldehyde for 48 hours at 4°C and decalcified in 10% EDTA for 14 days at 4°C. The samples were then dehydrated and embedded in paraffin for coronal sectioning using standard histology procedures. Paraffin tissue blocks were then sectioned at a thickness of 8 µm. Joint sections were then deparaffinized, rehydrated, and stained with Hematoxylin, Fast Green, and Safranin-O for histological grading. Two experienced graders (A.S. and S.Z.) evaluated the sections. Slides were randomized and assigned a temporary identification code to blind graders to diet treatment. OARSI mouse OA grading scores(*4*) and Mankin Scores(*5*) were assigned separately for the lateral femur, lateral tibia, medial femur, and medial tibia independently by each grader. Scores that differed by >2 between graders were re-evaluated for consensus scoring. Scores from the four compartments were then averaged to generate the average OARSI and Mankin scores.

### Western blotting

Mouse cartilage was dissected and collected in RIPA buffer and subjected to western blot analysis. Briefly, we first homogenized the cartilage tissue using a Scientific Industries Bead Genie Bead Beater (SI-B100). The supernatant that included proteins were subjected to electrophoresis using SDS-PAGE gels, transferred to PVDF membranes, and blocked for 30 minutes with 5% non-fat milk made in 1X TBST. The membranes were then incubated overnight with primary antibodies specific for malonyl-lysine (Cell Signaling Technology, #14942, 1:500 dilution) or anti-SIRT5 (Cell Signaling Technology, #8782, 1:500 dilution) diluted in TBS blocking buffer supplemented with 0.1% Tween-20. The membranes were then incubated with IRDye 800CW goat anti-rabbit secondary antibody (926-32211, 1:15,000 dilution). The membrane was imaged by Odyssey CLx Imaging System. Quantification of band intensity was performed in ImageJ.

### Proteomics data generation and analysis

Mouse primary chondrocytes (∼50 million cells) from WT or *Sirt5^-/-^*mice (n = 4 each) were homogenized in a solution containing 8 M urea, 200 mM triethylammonium bicarbonate (TEAB), pH 8, 75 mM sodium chloride, 1 μM trichostatin A, 3 mM nicotinamide, and 1x protease/phosphatase inhibitor cocktail (Thermo Fisher Scientific, Waltham, MA), and sonicated 2 times for 15 s on 30% amplitude using a probe sonicator. Lysates were clarified by spinning at 15,700 x g for 15 min at 4°C, and the supernatant containing the soluble proteins was collected. The extracts were then subjected to proteomics data generation as described before (*6*). Briefly, after protein concentration determination using Bicinchoninic Acid Protein (BCA) Assay (Thermo Fisher Scientific, Waltham, MA), protein samples were reduced, alkylated, and digested in solution with trypsin followed by peptide desalting with Oasis 30-mg Sorbent Cartridges (Waters, Milford, MA). Digest samples were vacuum dried and resuspended in 0.2% formic acid (FA) in water at a final concentration of 1 µg/µL. Finally, indexed retention time standard peptides (iRT; Biognosys, Schlieren, Switzerland) (*7*) were spiked in the samples according to manufacturer’s instructions. LC-MS/MS analyses were performed on a Dionex UltiMate 3000 system online coupled to an Orbitrap Eclipse Tribrid mass spectrometer (Thermo Fisher Scientific, San Jose, CA). The solvent system consisted of 2% ACN, 0.1% FA in water (solvent A) and 98% ACN, 0.1% FA in water (solvent B). Proteolytic peptides (1 µg) were loaded onto an Acclaim PepMap 100 C_18_ trap column (0.1 x 20 mm, 5 µm particle size; Thermo Fisher Scientific) over 5 min at 5 µL/min with 100% solvent A. Peptides were eluted on an Acclaim PepMap 100 C_18_ analytical column (75 µm x 50 cm, 3 µm particle size; Thermo Fisher Scientific) at 0.3 µL/min using the following gradient of solvent B: 2% for 5 min, linear from 2% to 20% in 125 min, linear from 20% to 32% in 40 min, up to 80% in 1 min, 80% for 9 min, and down to 2% in 1 min. The column was equilibrated at 2% for 29 min (total gradient length = 210 min). All samples were acquired in data-independent acquisition (DIA) mode (*8–10*). Briefly, full MS spectra were collected at 120,000 resolution (AGC target: 3e6 ions, maximum injection time: 60 ms, 350-1,650 m/z), and MS2 spectra at 30,000 resolution (AGC target: 3e6 ions, maximum injection time: Auto, NCE: 27, fixed first mass 200 m/z). The DIA precursor ion isolation scheme consisted of 26 variable windows covering the 350-1,650 m/z mass range with an overlap of 1 m/z (Suppl. Table 1. DIA_Isolation_Scheme) (*8*). DIA data processing and statistical analysis were performed in Spectronaut (version 15.1.210713.50606; Biognosys) using directDIA. Data was searched against all *Mus musculus* protein entries extracted from UniProtKB-TrEMBL (86,521 entries, 08/24/2021). Trypsin/P was set as digestion enzyme and two missed cleavages were allowed. Cysteine carbamidomethylation was set as fixed modification, and methionine oxidation and protein N-terminus acetylation as variable modifications. Data extraction parameters were set as dynamic and non-linear iRT calibration with precision iRT was selected. Identification was performed using 1% precursor and protein q-value. Quantification was based on the extracted ion chromatograms (XICs) of 3 – 6 MS2 fragment ions, and local normalization was applied. iRT profiling was selected. Protein groups were required to have at least two unique peptides.

Protein abundances were log2 transformed and quantile normalized, after which empirical Bayes-moderated *t*-tests were applied using the limma R package (*11*) to determine differential regulation in *Sirt5^-/-^* vs. WT samples. The Benjamini-Hochberg method was used to correct for multiple testing, with proteins with an adjusted *P* < 0.05 defined as being differentially regulated. Significantly upregulated and downregulated protein lists were then separately input into the Online Search Tool for Retrieval of Interacting Genes/Proteins (STRING, v11.5 (*12*)) to infer protein-protein interactions, with a default confidence score threshold of > 0.4 used to define interactions (as calculated based on a composite of all active interaction sources except for text-mining). In each case, sub-networks of interacting proteins were established using the Markov Cluster Algorithm with default inflation parameter of 3. Sub-networks containing ≥ 10 proteins were prioritized and further examined for (i) enrichment of Gene Ontology (GO) biological process terms and Reactome pathway terms, and (ii) hub protein classification. Enrichment analyses was carried out using the clusterProfiler (*13*) and ReactomePA R packages (*14*) as appropriate, with input background being all proteins tested for differential regulation, and enriched terms defined as those with a Benjamini-Hochberg corrected *P* < 0.05 enriched for at least 2 features. Hub protein status was characterized by eigenvector centrality score.

### Identification of the malonylome in chondrocytes using LC-MS/MS

LC-MS/MS to identify the malonylome was conducted in collaboration with the Cell Signaling Technology, Inc. Proteomics facility. Cellular extracts from WT and *Sirt5^-/-^* primary chondrocytes were sonicated, centrifuged, reduced with DTT, and alkylated with iodoacetamide.

7 mg total protein for each sample was digested with trypsin, purified over C18 columns (Waters) and enriched with the Malonyl-Lysine Motif Antibody (CST #93872) as previously described(*15*). LC-MS/MS analysis was performed using an Orbitrap-Fusion Lumos Tribrid Mass spectrometer as previously described (*15, 16*) with replicate injections of each sample. Briefly, peptides were separated using a 50cm x 100µM PicoFrit capillary column packed with C18 reversed-phase resin and eluted with a 90-minute linear gradient of acetonitrile in 0.125% formic acid delivered at 280 nL/min using an HCD-MS2 method. Full MS parameter settings are available upon request.

MS spectra were evaluated by Cell Signaling Technology using Comet and the GFY-Core platform (Harvard University) (*17–20*). Searches were performed against the most recent update of the NCBI *Mus musculus* database with a mass accuracy of +/-20 ppm for precursor ions and 0.02 Da for product ions. Results were filtered to a 1% peptide-level FDR with mass accuracy +/-5ppm on precursor ions and presence of a malonylated residue. All quantitative results were generated using Skyline (*21*) to extract the integrated peak area of the corresponding peptide assignments. Accuracy of quantitative data was ensured by manual review in Skyline or in the ion chromatogram files. A complete list of identified peptides and proteins can be found in Suppl. Table 2_OhioU_00019005_MalK_FINAL. KEGG pathway enrichment analysis for proteins with ≥ 1 MaK site was performed via the Database for Annotation, Visualization and Integrated Discovery (*22*) using the Uniprot reviewed mouse proteome as background, with significantly enriched pathways defined as those with a Benjamini-Hochberg corrected P-value < 0.05 enriched for at least 2 features.

### Bulk RNA-Sequencing

WT and *Sirt5^F101L^* transfected primary chondrocytes were used for bulk RNA-seq analysis. RNA extraction, quality control, and RNA-sequencing (RNA-seq) were performed by Novogene. Data analysis was performed in-house as previously described (*23*).

### Measurement of GAPDH activity

The activity of Glyceraldehyde 3-phosphate dehydrogenase (GAPDH) was quantified using a GAPDH activity assay kit (Abcam, ab204732). Primary immature chondrocytes were extracted from the knee cartilage of 7-day-old wild-type mice and cultured in DMEM medium supplemented with 10% fetal bovine serum and 1% penicillin/streptomycin solution. At lower passages, these cells were transfected with plasmids encoding either wild-type GAPDH or the GAPDH K184E mutant (with a control group receiving no DNA). The efficiency of transfection was confirmed via western blot analysis. After 48 hours of transfection, cells were harvested by trypsinization, washed with 1X PBS, and resuspended in assay buffer at a concentration of 1 x 10^6 cells per 100 µL. The cells were homogenized by passing the suspension through a 25G syringe (1 mL) 10-15 times. The homogenized suspension was then incubated on ice for 15 minutes, followed by centrifugation at 10,000g for 5 minutes at 4°C. An aliquot of 25 µL of the supernatant was used to measure enzyme activity according to the manufacturer’s protocol. Absorbance was measured at OD 450 nm in kinetic mode using a BioTek microplate reader. A total of 30 readings were recorded over 30 minutes for three technical replicates of each sample. GAPDH activity was calculated based on the manufacturer’s guidelines.

### Lactate assay

The amount of lactate produced in the samples was quantified using the L-lactate colorimetric assay kit (Abcam, ab65331). Primary immature chondrocytes at lower passages were transfected with wild-type GAPDH or the GAPDH K184E mutant (with a control group receiving no DNA). After 48 h of transfection cells, 2 X 10^6^ cells/sample were harvested and resuspended in 4X volume of lactate assay buffer after thoroughly washing with 1X PBS. Cells were homogenized quickly passing through a 25G syringe several times. Cell homogenate was then centrifuged for 5 mins at 4°C at top speed to remove insoluble material. The supernatant was collected in a fresh tube and subjected to PCA/KOH deproteinization step according to the manufacturer’s protocol to remove the endogenous lactate dehydrogenase enzyme that will degrade lactate. An aliquot of 50 µL of the samples was used for quantifying the amount of lactate according to the manufacturer’s protocol. Colorimetric absorbance from the reaction was measured at OD 450 nm using a BioTek microplate reader. The concentration of L-lactate in the sample is calculated as:

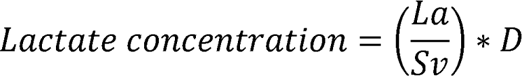

where: La = amount of lactic acid in the sample well calculated from a standard curve (nmol).

Sv = volume of sample added in the well (µL).

D = sample dilution factor.

### Diagnostic and procedure codes used to identify individuals with osteoarthritis in the Utah Population Database (UPDB)

Our coding strategy used to identify individuals with erosive hand, 1^st^ MTP joint, distal and proximal interphalangeal joint, thumb joint, and shoulder osteoarthritis (OA) has been previously described in detail (*23–25*). Briefly, the following diagnostic codes (ICD-9 and ICD-10) and procedure codes (CPT - Current Procedural Terminology) were used to identify affected individuals in the Utah Population Database.

<u>a)</u> Distal and proximal interphalangeal joint OA:

CPT: 26862, 26863, 26860, 26861 (arthrodesis, interphalangeal joint, with or without internal fixation) and 26535, 26536 (arthroplasty, interphalangeal joint).

ICD-9: 715.14 (osteoarthritis, localized, primary, hand).

ICD-10: M19.04x (primary osteoarthritis, hand).

<u>b)</u> 1^st^ MTP joint OA (synonymous with hallux rigidus):

CPT: 28289 (hallux rigidus correction with cheilectomy, debridement and capsular release 1^st^ metatarsophalangeal joint) and 28750 (arthrodesis, great toe).

ICD-9: 735.2 (hallux rigidus). ICD-10: not used.

<u>c)</u> Erosive hand OA:

ICD-10 M15.4 (Erosive (osteo)arthritis).

<u>d)</u> Glenohumeral OA:

CPT: 23472 (arthroplasty, glenohumeral joint).

ICD-9: 715.11 (osteoarthritis, localized, primary, shoulder region).

ICD-10: M19.011 (primary osteoarthritis, right shoulder), M19.012 (primary osteoarthritis, left shoulder), M19.019 (primary osteoarthritis, unspecified shoulder), Z96.611 (presence of right artificial shoulder joint), Z96.612 (presence of left artificial shoulder joint), or M19.0x (primary osteoarthritis, shoulder).

<u>e)</u> Exclusion criteria:

Individuals diagnosed with any of the following codes were excluded:

ICD-9: 714.0 (rheumatoid arthritis), 714.2 and 714.3 (rheumatoid arthritis and other inflammatory polyarthropathies).

ICD-10: M05.xxx (rheumatoid polyneuropathy with rheumatoid arthritis), M06.xx (other rheumatoid arthritis), or M08.xxx (juvenile arthritis).

were excluded if they were diagnosed with any of the following codes:

ICD-9 codes: ICD-9 716·13, 716·14 (traumatic arthropathy, forearm and hand, respectively). ICD-10 codes: M18·2x, M18·3x (post-traumatic), M18·4 (‘other’ etiology), M18·5x (‘other’ etiology), M18·9 (unspecified etiology)

Inflammatory arthritis codes: ICD9 714·0, 714·2, 714·3, ICD10 M05·xxx, M06·xx, M08·xxx. Ligamentous hyperlaxity codes: 728·4 (laxity of ligament), 718·84 (hypermobility/instability of joints, including hand), 756·83 (Ehlers-Danlos syndrome), M35·7 (hypermobility syndrome), M24·2x (ligamentous laxity NOS), Q79·6x (Ehlers-Danlos syndrome).

Individuals were asked if they were diagnosed with psoriatic arthritis, gout, or had a traumatic injury to the affected joint. If they answered yes, they were excluded from the study. In these high-risk, multigenerational families, the frequency of OA-affected members is significantly higher than expected based on UPDB analysis as a whole. Although these families were initially identified based on joint-specific phenotypes, detailed chart reviews revealed that individuals often have OA in other joints as well (e.g., knee and hip OA in a family primarily identified with hand OA). The pathways we discover are likely shared by most or all synovial joints, with additional cofactors influencing the probability of a particular joint being affected.

### Identification and sequence analysis of families with inherited OA

Families with a statistically significant enrichment of OA that appeared to segregate as a simple dominant Mendelian trait were identified from the UPDB OA cohort (*24–26*). Selected members were subjected to whole exome sequencing and candidate causative variants were identified (*24, 26*). Briefly, whole exome sequencing (WES) and analysis was performed using genomic DNA isolated from whole blood or saliva as previously described (*23*) (*24, 26*). Libraries were prepared using the Agilent SureSelect XT Human All Exon + UTR (v8) kit followed by Illumina NovaSeq 6000 150 cycle paired end sequencing. We followed best practices established by the Broad Institute GATK for variant discovery (https://gatk.broadinstitute.org/hc/en-us). Analysis of variants was performed with ANNOVAR (*27*) (http://annovar.openbioinformatics.org/en/latest/) and pVAAST (*28*) (http://www.hufflab.org/software/pvaast/) in concert with PHEVOR2 (http://weatherby.genetics.utah.edu/phevor2/index.html).

## Data and materials availability

### Proteomics data availability

Raw data and complete MS data sets have been uploaded to the MassIVE repository of the Center for Computational Mass Spectrometry at UCSD, and can be downloaded using the following link: https://massive.ucsd.edu/ProteoSAFe/dataset.jsp?task=698e43b08c354177bb5227ac7cc400f5 (MassIVE ID: MSV000095237; ProteomeXchange ID: PXD053614).

[Note to the reviewers: To access the data repository MassIVE (UCSD) for MS data, please use: Username: MSV000095237_reviewer; Password: winter].

<u>RNA-Seq data availability</u>: RNA-Seq raw data has been uploaded to the Gene Expression Omnibus database: GSE272422 (https://www.ncbi.nlm.nih.gov/geo/query/acc.cgi?acc=GSE272422)

Malonylome data availability: Raw data and complete MS data sets associated with the chondrocyte malonylome have been uploaded to Google Drive: https://drive.google.com/drive/u/1/folders/15aHZFb-bCEJ-1LoU8F9NpM-YbYNDcQTs

## Notes

### Competing Interest Statement

The authors have declared no competing interest.

### Summary of Updates

We updated the authors as well as uploaded the supplementary methods.

